# Distributional reinforcement learning in prefrontal cortex

**DOI:** 10.1101/2021.06.14.448422

**Authors:** Timothy H. Muller, James L. Butler, Sebastijan Veselic, Bruno Miranda, Timothy E.J. Behrens, Zeb Kurth-Nelson, Steven W. Kennerley

## Abstract

Prefrontal cortex is crucial for learning and decision-making. Classic reinforcement learning (RL) theories centre on learning the expectation of potential rewarding outcomes and explain a wealth of neural data in prefrontal cortex. Distributional RL, on the other hand, learns the full distribution of rewarding outcomes and better explains dopamine responses. Here we show distributional RL also better explains prefrontal cortical responses, suggesting it is a ubiquitous mechanism for reward-guided learning.

## Main text

The prefrontal cortex (PFC) is critical for appropriate learning and decision-making^1–6^. Reinforcement learning (RL) theory offers a computational framework for understanding learning and decision-making processes^7^ and accordingly explains many neural responses throughout PFC^8,9^. “Classic” RL models^7,10^ learn to predict the expectation – or mean – of the distribution over possible rewarding outcomes following a stimulus or action. However, by learning only the expected reward, knowledge of the underlying reward distribution – which is essential for risk-sensitive decision-making – is lost. Furthermore, since in classic RL models all neurons learn to predict the same, expected reward, the classic RL framework is unable to account for substantial diversity in reward-related responses across PFC neurons^8,11,12^.

A recent modification to classic RL – distributional RL – learns the full reward distribution and offers a candidate explanation for neuronal diversity^13–15^. Unlike classic RL models, in distributional RL, different neurons learn to predict different parts of the reward distribution. Some neurons will carry value predictions above the mean of the reward distribution, and others below – referred to as optimistic and pessimistic neurons, respectively. This means that across the population of neurons the full distribution of possible rewards is encoded, and that diversity is predicted in neuronal responses. By explaining such diversity, recent research demonstrates distributional RL better explains responses of midbrain dopaminergic neurons^15^ – famously known to encode reward prediction errors (RPEs) that drive learning of reward predictions^16^ – than classic RL.

The PFC encodes a diversity of learning- and decision-related computations^8,17,18^, including reward prediction errors^8,19^. However, a striking feature of PFC, particularly the anterior cingulate cortex (ACC), is that it encodes diverse reward-related features, such as diverse temporal scales and learning rates^8,20–22^, suggesting PFC neurons may be a strong candidate for distributional RL. Furthermore, PFC is engaged in risk-sensitive decision-making tasks^23^, for which distributional representations will be useful. Given the findings of distributional RL in dopamine neurons, and the fact that PFC is a major recipient of dopaminergic input^24–26^, we examined whether distributional RL explains reward responses in multiple regions of primate PFC in two different decision-making tasks. In the first dataset, we demonstrate key signatures of distributional RL conceptually analogous to those shown in mouse dopamine neurons^15^. In the second dataset, we provide evidence for a previously untested prediction of distributional RL; that there are asymmetries in the rates of learning from better vs. worse than expected outcomes.

To test for signatures of distributional RL in the first dataset, we tested three key predictions. The first is that different neurons carry different value predictions, varying in their level of optimism. In contrast to classic RL, which predicts all neurons linearly increase their firing rate as a function of reward and no diversity across neurons (since all neurons predict the expected reward), the varying value predictions in distributional RL manifest as: i) neurons increasing their firing rate non-linearly as a function of reward, and ii) diversity in this non-linearity across neurons^15^ (Figure 1B). The same value-predicting cue will therefore elicit different responses in different neurons. The second prediction of distributional RL is that different neurons will place different relative weights on positive vs. negative RPEs. Because the different value predictions arise from these different relative weights, the third prediction of distributional RL is that these two sources of diversity should correlate.

**Figure 1.**
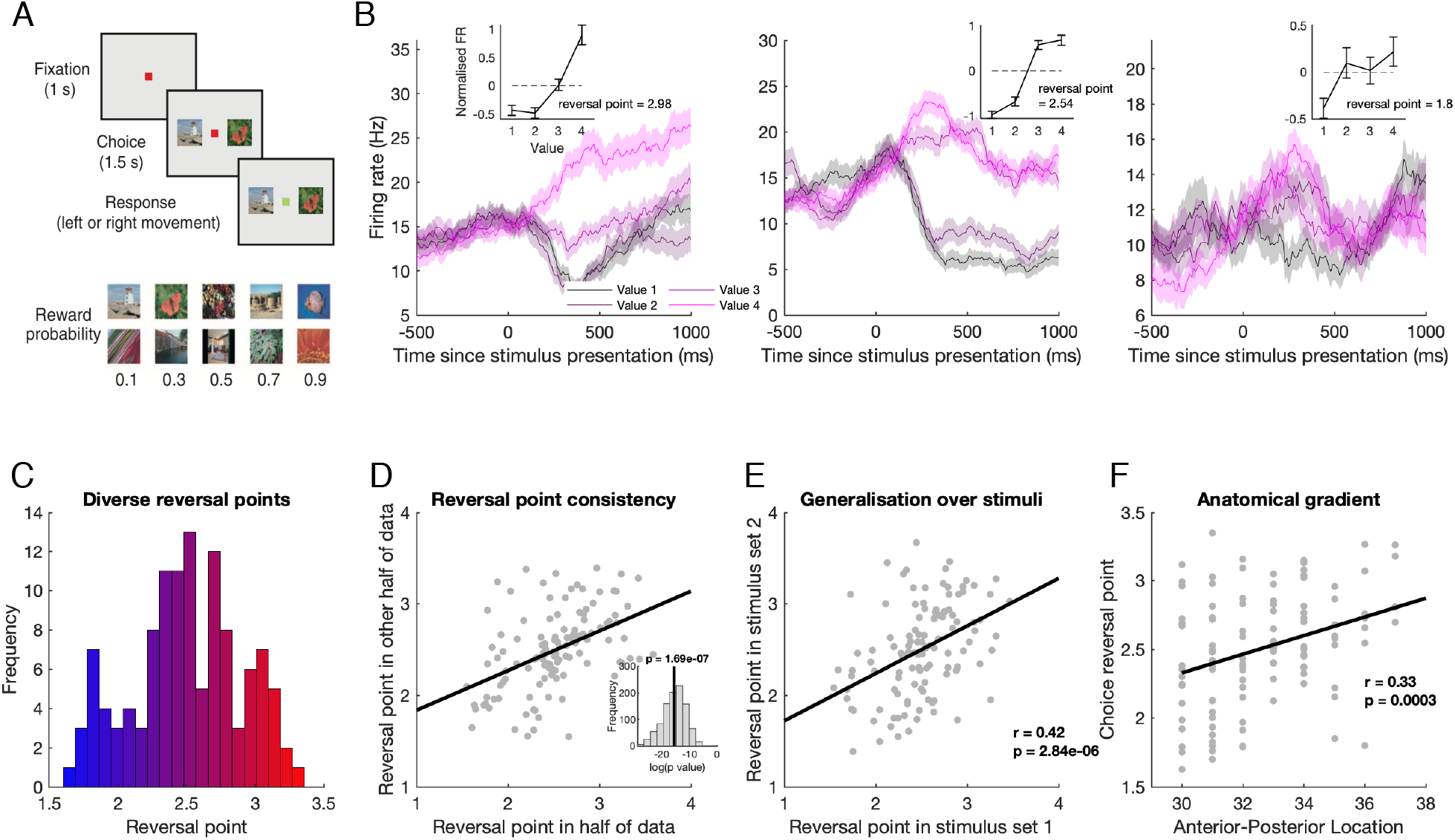
Diverse optimism in value coding across ACC neurons. A) On each trial, subjects chose between two cues of neighbouring probability value. Each probability value could be denoted by two stimuli, resulting in two stimulus sets (see^8^ for task details). B) Example neuron responses demonstrating different levels of optimism. In each plot the mean firing rate is plotted as a function of time and split according to the chosen value (probability) level. There are four chosen values (0.3-0.9 probability) as subjects rarely chose the 0.1 probability level (choice accuracy was at ceiling; 98%). Insets demonstrate firing rate is a non-linear function of value. Mean firing rate (having z-scored across trials all trials) in a 200-600ms window post-cue onset is plotted as a function of the 4 values. Reversal points are the interpolated value at which there is 0 change from the mean firing rate, an index of non-linearity. C) Histogram showing a diversity of reversal points across neurons. D) Scatter plot showing reversal points estimated in one half of the data strongly predict that in the other half. Each point denotes a neuron. Inset: log p-values of the correlation between 1000 different random splits of the data into independent partitions. Black line denotes the geometric mean of these p-values. E) Scatter plot showing reversal points estimated in stimulus set 1 strongly predict those in stimulus set 2. Each point denotes a neuron. F) Anterior-posterior topographic location of the neuron predicts its reversal point, with more anterior ACC neurons more optimistic. Each point denotes a neuron.

To test the first prediction of distributional RL, we examined non-linear value coding in neurons recorded from three PFC regions implicated in learning and decision-making^8,9,18^; the lateral prefrontal cortex (LPFC, n=257), the orbitofrontal cortex (OFC, n=140), and the anterior cingulate cortex (ACC, n=213). Two non-human primates (NHPs, *Macaca mulatta*) were presented with a choice of two value-predicting stimuli, which varied in the probability of upcoming reward while magnitude of reward was held constant^8^ (Figure 1A). We initially analysed firing rates of reward-sensitive neurons (Supplementary Information) as a function of the 4 possible chosen value (probability) levels by looking at the mean firing rate in a window of 200-600ms post-stimulus onset during the choice epoch of the task. We then subtracted the mean firing rate in this window across trials. We observed non-linearities in individual neurons, which we indexed using a measure of optimism analogous to Dabney et al.^15^. We measure the ‘reversal point’ of a given neuron using the mean-subtracted firing rates to find the interpolated cue value for which the firing rate reverses from above to below the mean firing rate (Figure 1B). In classic RL, we would expect the reversal point for all neurons to be around the mean of the value distribution (i.e., around 2.5 in this dataset), whereas optimistic neurons in distributional RL will have reversal points falling towards higher values, and vice versa for pessimistic neurons (Figure 1B).

We observed diversity in reversal points (i.e., value predictions) across the population of 117 reward-sensitive neurons in ACC, which exhibited both optimistic and pessimistic neurons (Figure 1B and 1C). To confirm this diversity in ACC is not simply due to noise, we demonstrated that the reversal point estimated in a random half of the data predicted that in the other half of the data (*R* = 0.46, *P* = 1.7×10^−7^ by Pearson correlation; Figure 1D). Neurons in OFC and LPFC exhibited lower reward-selectivity, and a lack of diversity in reversal point (Supplementary Information), and hence we focus on ACC for the remainder of this report.

Critically, distributional RL predicts reversal diversity is a signature of distributional coding over value, not over stimulus features. A neuron tuned to the sensory (e.g. visual) features for the cue predicting value 4 would appear as an optimistic neuron by distributional RL definitions, even though it may not be optimistic. Our experiment controlled for this by using two stimuli for each value level. We showed optimism in ACC generalises over different stimulus sets by correlating the reversal point estimated in one stimulus set with that of the other (*R* = 0.41, *P* = 2.8×10^−6^ by Pearson correlation; Figure 1E). This confirmed that diversity in ACC reversal points is neither explained by noise nor tuning to specific stimulus features.

In RL settings, ACC exhibits a spectrum of learning rates topographically organized along the posterior-anterior axis^22^. We therefore tested for topographic organisation of optimism in ACC reward-selective neurons, and found the anterior-posterior location within ACC predicted optimism, such that more anterior neurons were more optimistic (*R* = 0.33, *P* = 2.9×10^−4^ by Pearson correlation; Figure 1F). Such topographic organisation to optimism may ensure neurons interact with other neurons of similar optimism^15,27^, and may offer a route to non-invasive measurement and manipulation of optimism.

A second key prediction of Distributional RL is that different value predictions arise from different relative weights placed on positive vs. negative RPEs^15^. If, for example, a neuron puts more weight on – and therefore learns more from – positive than negative RPEs, it will learn an optimistic value prediction. Asymmetries in the relative weighting of positive and negative RPEs can be indexed by the relative scaling, i.e. regression slope, of positive and negative RPEs (Figure 2A). For each neuron, we determined separately the scaling of positive and negative RPEs elicited at feedback. We did this by regressing firing rate against the chosen probability cue – which determines the size of the RPE – on rewarded and unrewarded trials, respectively. From these we computed a single measure to reflect the asymmetry of positive vs. negative RPEs; *β*^+^/(*β*^+^ + *β*^−^), where *β*^+^ and *β*^−^ are the betas for positive and negative RPEs, respectively. We include in this analysis only those neurons that are RPE selective (41 neurons in ACC, see Supplementary Information). We found diversity in the relative weighting of positive vs. negative RPEs across ACC neurons at feedback (Figure 2B), and this diversity was stable across independent partitions of the data (*R* = 0.32; *P* = 0.015 by Pearson correlation; Figure 2C).

**Figure 2.**
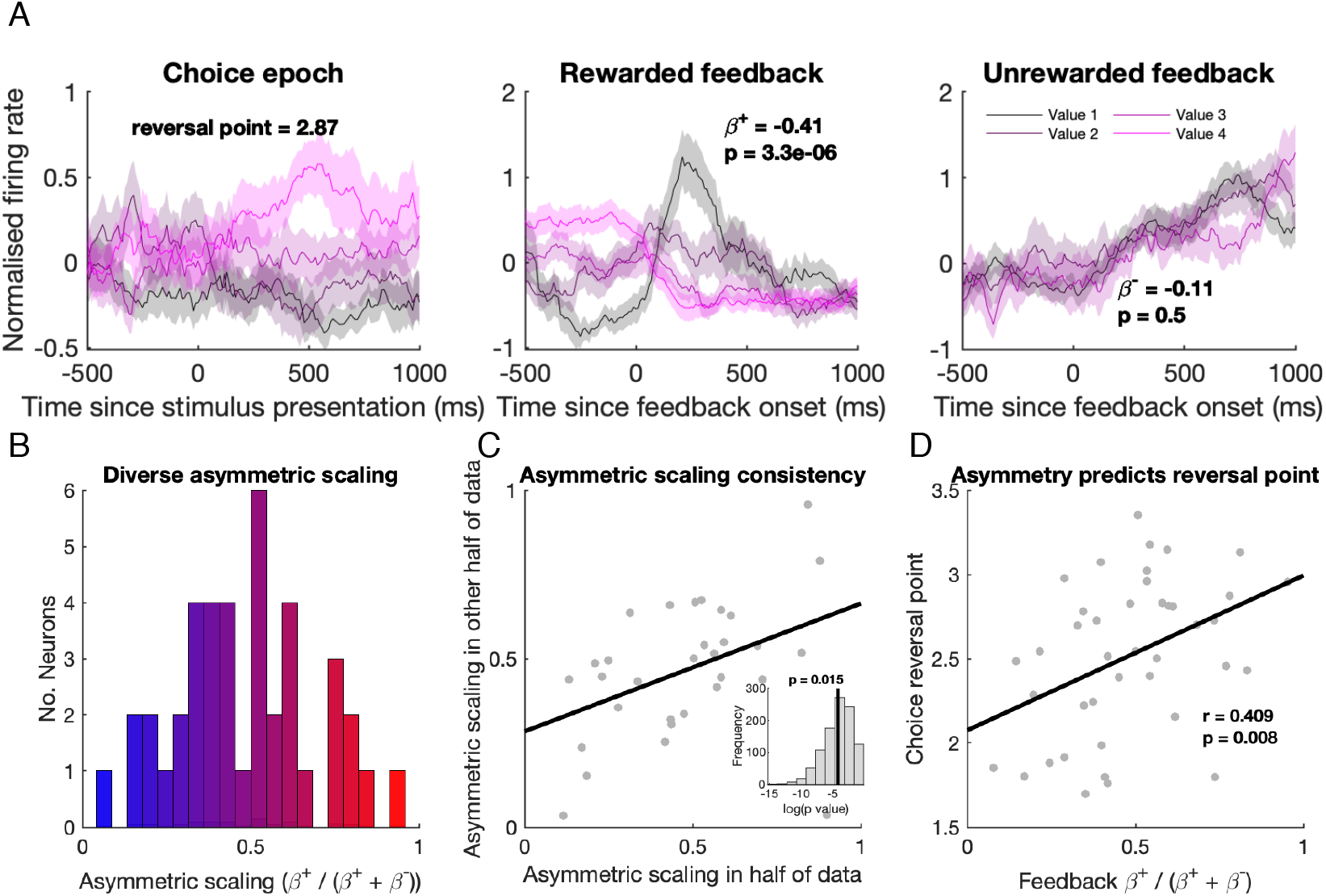
Diverse asymmetric scaling of reward prediction errors predicts choice optimism in ACC. A) An example neuron’s responses at each of the task epochs: choice, feedback on rewarded trials, and feedback on unrewarded trials, demonstrating where in the task the betas are measured for RPE scaling asymmetries. *β*^+^ and *β*^−^ are betas corresponding to the scaling of positive and negative RPEs. B) Histogram showing a diversity of asymmetric scaling across ACC RPE neurons. C) Same format as Figure 1D but for asymmetric scaling consistency. Each point denotes a neuron; RPE-selective neurons only. D) Asymmetric scaling estimated at feedback predicts reversal point at choice. Each point denotes a neuron.

While we know neuronal responses in cortex are diverse, often talked about in the context of mixed selectivity^11,28,29^, distributional RL makes a more specific prediction. Since, in distributional RL theory, the different weights placed on positive vs. negative RPEs result in the diverse levels of optimism in value predictions that we observed at choice, these should be related. This is despite being two different measures of optimism estimated in different periods of the task (i.e., choice and outcome). We confirm this prediction in ACC neurons; asymmetry in RPEs at feedback predicted the reversal points at choice (*R* = 0.41, *P* = 0.0079 by Pearson correlation; Figure 2D). Thus primate PFC contains conceptual analogues of distributional RL found in rodent dopamine neurons^15^.

Thus far we have identified neural signatures of distributional RL in PFC during decisions where the reward structure is static and values do not need to be updated through learning. However, many real world contexts require continuous learning as decision values change. We now turn to a previously untested, strong prediction of distributional RL, which predicts a unique pattern of learning across neurons. That is, in addition to diverse asymmetries in the *scaling* of positive vs. negative RPEs, distributional RL also predicts diverse asymmetries in the rates of *learning* from positive vs. negative RPEs (Figure 3A and 3B).

**Figure 3.**
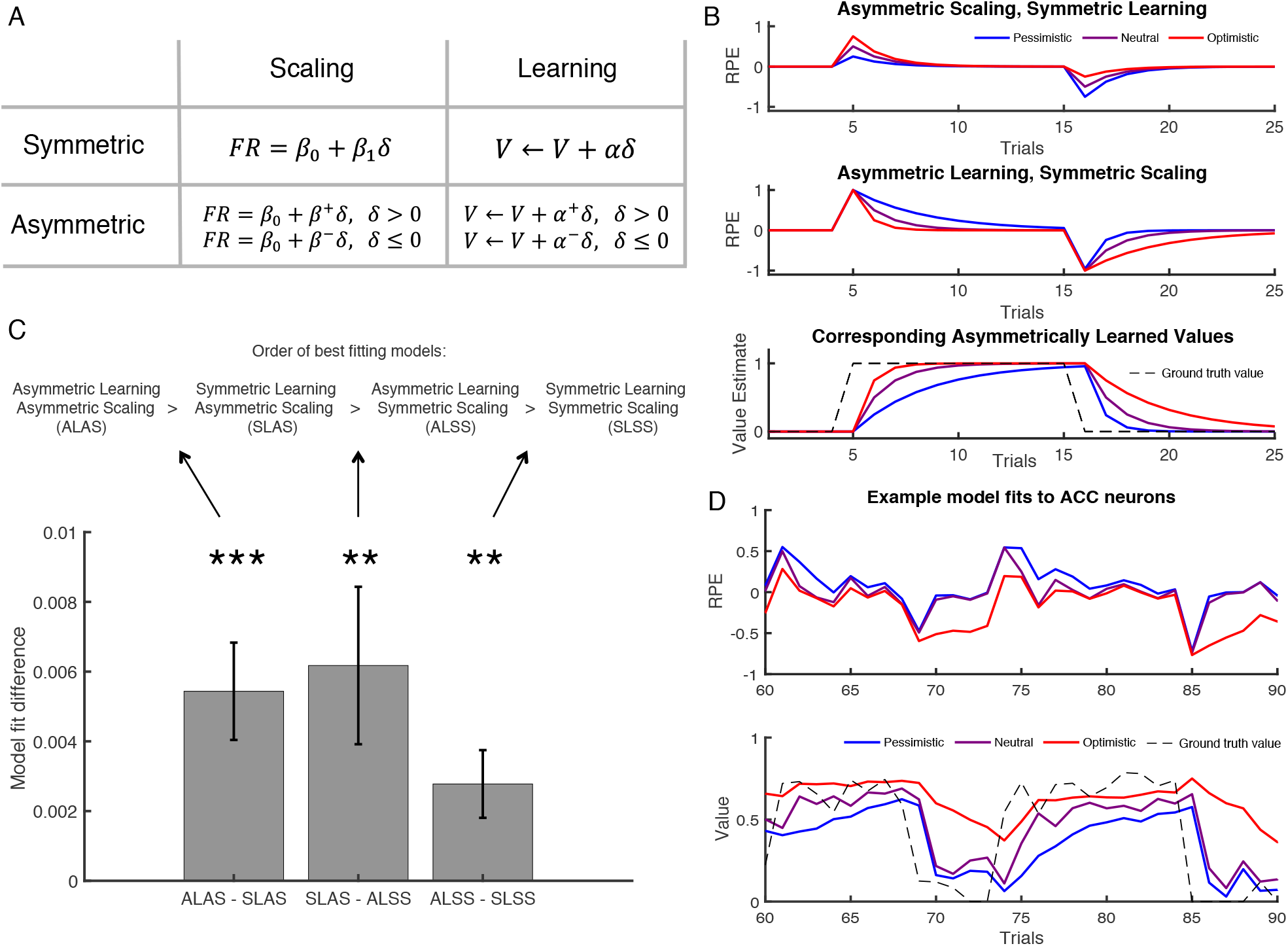
Asymmetric learning. A) Asymmetric scaling and asymmetric learning are both predictions of distributional RL, but dissociable. Asymmetric scaling reflects differences in the degree to which positive and negative RPEs are scaled in order to predict firing rate. Asymmetric learning reflects differences in the rate of state value update following positive and negative RPEs. These different learning rates are denoted by *α*^+^ and *α*^−^, respectively. *δ* = *r* − *V* is the RPE, where *r* is the reward on the current trial and *V* the value. B) Simulated examples demonstrating the difference between asymmetric scaling and learning. C) Comparing cross-validated model fits reveals that a model with both asymmetric scaling and asymmetric learning is the best explanation of the ACC data. This is followed by a model with asymmetries in only scaling, then only learning, and the fully classic (symmetric) model is the worst model of the data. Each bar in the bar graph shows the comparison between a pair of models. D) Example model fits. Top: RPE regressors generated using learning rate parameters fit to individual neuron data, for three different neurons from the same session. Different levels of optimism can be seen via the different rates at which RPEs tend back toward zero following changes in state value (denoted by dashed black line in bottom plot). Bottom: This is reflected in the corresponding values. The pessimistic neuron (shown in blue), for example, is quick to devalue but slow to value.

Whereas in *asymmetric scaling*, neurons differ in the degree to which the firing rate response is scaled as a function of positive vs. negative RPEs, in *asymmetric learning*, the rate at which a neuron updates its estimated value of a state from RPEs differs for positive vs. negative RPEs (Figure 3A). This will influence subsequent RPE responses as these are computed from the state value. For example, an optimistic neuron will increase its predicted value more rapidly than decrease it following changes in the true value. Therefore, following an increase in the true value, the size of positive RPE responses will decrease more rapidly since the new (higher) reward is learnt – and therefore expected – more rapidly. The converse pattern will be true for pessimistic neurons (Figure 3B).

Exploring asymmetric learning requires a learning task in which the reward structure is dynamic, so that subjects must update their value expectations. It also requires neurons that are outcome selective. We recorded single neuron data from two different NHPs (*Macaca mulatta*) in ACC during performance of the “two-step task”^30,31^, in which there were 4 end-stage cues that independently changed in value every 5-9 trials (Supplementary Information). Subjects were therefore required to update their value estimates of these cues across trials. We identified a significant population of reward selective neurons in ACC (n=111/240, 46%), which were then used for the asymmetric learning analysis below (Supplementary Information).

To test for asymmetric learning, we modelled neuronal responses using classic (temporal difference RL, which learns the expected value) and distributional RL models and tested which best fits the data (Supplementary Information for details). We adapt the one-step transition temporal difference RL model to incorporate either asymmetries in scaling, asymmetries in learning, or both, and asked which model best explained the data. For each neuron, we fit the learning rate and scaling parameters in a subset of the data. We then used these parameters to generate RPE regressors for the held-out data in which we assessed the model’s fit to the data (R-squared) using 10-fold cross validation. We then compared the different models using the mean R-squared (across partitions) for each neuron.

We found a model that incorporates both asymmetric learning as well as asymmetric scaling was the best fit to the ACC data (Figure 3C). This suggests that two learning rates are better than one in explaining the learning dynamics of ACC neuronal responses; demonstrating that different neurons update their value estimates at different rates, depending on whether they are increasing vs. decreasing in value. For example, one neuron may increase its value estimate rapidly for positive outcomes, but decrease it slowly for negative outcomes, and vice versa for other neurons (Figure 3D). This finding in ACC provides evidence for a previously untested key prediction of the distributional RL theory.

Distributional RL provides a powerful new computational framework that learns the full reward distribution rather than only the expectation, improves performance of artificial agents, and better explains rodent dopaminergic responses^13–15^. Here, we provide the first demonstration that distributional RL also better explains single-neuron responses in cortex, specifically in primate ACC. We show conceptual analogues to the results in dopamine; specifically, evidence for diverse value predictions that are correlated with diverse asymmetries in RPE scaling. We further show diversity generalises over stimulus features, demonstrating it is due to diverse optimism not stimulus feature coding, and that it lies on an anatomical gradient, offering a candidate solution to the connectivity patterns predicted by distributional RL^15,27^. Finally, we present evidence for asymmetries in the rates of learning from positive vs. negative RPEs.

That distributional RL is in cortex has several important implications. First, it provides a candidate mechanism for how cortical representations of probability distributions over value arise, as are required for value-based risk-sensitive decision-making^23,32^. Second, PFC responses are famously diverse, with different neurons showing different selectivity profiles^11,18,29^. Although there is much diversity still to be explained in PFC, distributional RL provides one account for such diversity and suggests other interesting computational principles may underlie other sources of diversity. Third, it raises intriguing questions about possible differences between dopaminergic and cortical distributional RL. Perhaps, the latter facilitates learning cortical representations^15,33^ that implement the former, which is used for risk-sensitive decision-making. Finally the presence of distributional RL in cortex of NHPs across two different studies suggests distributional RL may be hidden in many RL datasets, and that it may be a ubiquitous mechanism for reward-guided learning.

## Supplementary information

### Task and neural recordings from Kennerley et al 2011

Results in Figures 1 and 2 are a re-analysis of the data presented in Kennerley et al 2011^8^. Full task and recording details can be found there, but we outline the key points relevant to the current study here.

This task was a two-alternative forced choice task, in which two rhesus macaques were presented, on each trial, with two stimuli, which they chose between using a joystick movement. After a delay feedback was delivered. Trials differed in the pair of stimuli presented at the choice phase. Stimuli were drawn from a set of possible stimuli, which denote different values varying along one of three attributes: probability of reward, magnitude of reward, or the amount of effort (lever pulls) required to obtain reward.

On any given trial, subjects will be presented with two stimuli that: belong to the same attribute (e.g. they are both probability cues), will be drawn from the same stimulus set within that attribute, and denote neighbouring values (e.g. subjects will choose between 0.9 and 0.7 probability cues and never 0.9 and 0.5). This means animals only ever choose between options within the same attribute and stimulus set, and that the chosen value difference is the same on all trials. For the purpose of this study, our analyses only focus on probability (not magnitude or effort) trials.

Recordings were made in ACC (n=213 neurons), OFC (n=140) and lateral PFC (n=257); see figure 6 of Kennerley et al 2009^34^ for precise locations of recorded neurons.

This dataset is well suited to test for distributional RL given recordings were in ACC, a region known to contain value-related learning signals^8^ and to be important for risk-sensitive decision-making^23^. Furthermore, this dataset is well suited as we can index neural responses to positive and negative RPEs separately^8^ (see below). Indeed, we previously reported^8^ that some neurons in ACC encode, for example, positive RPEs more strongly than negative RPEs, which is suggestive – broadly speaking – of diversity in RPE coding. In addition to being the most appropriate brain region to test for distributional RL in cortex, the ACC is also recorded in both this dataset and the other dataset analysed in this manuscript (see below).

Note that we only present results from the analyses of probability trials, since these are the only trials that we can measure asymmetric scaling of RPEs because the probabilistic feedback causes positive and negative RPEs on rewarded and unrewarded trials, respectively (see below).

### Neuron inclusion criteria and analysis assumptions

Only neurons that significantly encoded reward probability at choice (“probability-selective”) entered into subsequent distributional RL analyses. Probability-selective neurons were defined as: P<0.05 in linear regression between probability level and mean firing rate on each trial in a 200-600ms window post-cue onset. This is the analysis window used throughout the paper. 117 ACC neurons (55%) passed this criterion. In contrast, only 30% of OFC and 30% of LPFC neurons met this criterion. For the asymmetric scaling analyses in Figure 2, we include the subset of these probability-selective neurons that additionally encode reward probability at feedback with an opposite sign to that at choice, i.e. reward prediction error (RPE) selective neurons (see below) as we defined previously^8^. 41 (19%) ACC neurons met this criterion; in contrast, only 6% of OFC and 10% of LPFC neurons met this criterion. Furthermore, there was a lack of significant diversity in reversal point in OFC and LPFC (Supplementary Figure 3). Hence we focused on the ACC for the remainder of the analyses.

We briefly note here that, unlike dopaminergic neurons, in ACC, some neurons’ firing rates have a positive relationship with reward (i.e., firing rate increases as reward increases) and others negative (i.e., firing rate increases as reward decreases)^8^. We therefore flipped the firing rates (multiply by −1) of those neurons that are negative, but note that this in fact does not make any difference to the estimation of the distributional RL measures.

Furthermore, since the value differences of the choices are constant (as they are only ever shown pairs of stimuli neighbouring in value) and their performance is at ceiling^8^ (choosing the higher option on 98% of trials), we have 4 possible values corresponding to the 4 possible pairs of stimuli that could be presented on each trial.

Note that some trials are better than others and therefore ACC responses to the choice cues could reflect an RPE of the current trial value relative to the average set of trial values that could be offered. We can therefore treat the response to the cues at choice as a RPE (i.e., current offer value – average trial value).

### Measuring optimism at choice

We index the non-linearity in the firing rate as a function of reward using a measure analogous to that used in Dabney et al 2020^15^. We measure the ‘reversal point’ of a given neuron by estimating the value at which that neuron’s response is the same as (or reverses from positive to negative deviation from) the mean firing rate across trials following the presentation of the value-predicting cue (in the analysis window).

We note that, unlike in dopaminergic neurons, the reversal point here is induced by z-scoring the data (mean firing rate in the analysis window post-stimulus onset) across trials, and is therefore not the same as reversal point from baseline (pre-stimulus onset) firing, as used in Dabney et al^15^. This is necessary because deviation in firing rate from baseline in cortical neurons does not have the same assumed meaning as it does in dopaminergic neurons. In dopaminergic neurons, it is assumed that positive and negative deviations from baseline firing rate equate to positive and negative RPEs being signalled by that neuron^15,16^. However, in cortex, many probability selective neurons will, for example, increase their firing rate (relative to the pre-cue baseline) in response to all values (i.e. even those at the lowest part of the reward distribution, which ought to elicit negative RPEs even in the most pessimistic neurons). Hence, unlike in dopaminergic neurons, in cortex, an increase in firing rate relative to baseline does not necessarily mean a positive RPE. We therefore measure the reversal point for all neurons by z-scoring the data in a window after feedback, so that we can compare the measures of optimism across neurons (this z-scoring results in neutral neurons having a reversal point of 2.5 and deviations above and below this indicate optimism and pessimism, respectively). If a neuron is optimistic and thus predicts the highest values in the range of the task, the firing rate to all values but the highest value will be low relative to that of the highest, hence the reversal point will be high (see Fig 1B, left). We use this reversal point measure for consistency with Dabney et al^15^.

However, we note that an alternative measure of optimism capturing the non-linear shape of the neuronal response as a function of reward gives qualitatively the same results (Supplementary Figure 1), and is highly correlated with the reversal point. This measure is obtained by fitting the non-linearity in the firing rate as a function of reward using a quadratic term in linear regression:

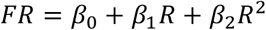

where *FR* is the firing rate on each trial and *R* the reward level. *β*_2_ is a regression weight that indexes optimism via the concavity (or convexity) of the function. As expected, this measure of optimism is highly correlated with the reversal point described above (*R* = 0.87, *P* = 4.0×10^−37^ by Pearson correlation) – corroborating that both measures index the non-linearity in the firing rate as a function of reward.

### Consistent diversity in optimism at choice

Observing diversity in optimism/reversal point alone is not sufficient, as this would be expected simply by noise. We therefore confirm diversity in reversal points is consistent by partitioning the data into independent partitions and testing whether the diversity is consistent across the partitions. We follow the same methodology as in Dabney et al 2020^15^. We estimate the reversal point in a random half of the trials, and repeat in the other half. We do this for each neuron and then correlate the reversal points estimated in one half with that in the other, obtaining r and P values for the correlation. If diversity is not due to random noise, we would expect these independently estimated reversal points to significantly correlate across neurons. To ensure this correlation is robust across partitions of the data, we repeat this partitioning process 1000 times and take the geometric mean of the P-values across partitions to obtain a summary P-value for the analysis.

### Asymmetric scaling (RPEs at feedback) analysis

In order to estimate asymmetry in the scaling of firing rate as a function of positive vs. negative RPEs, we estimated the scaling of positive and negative RPEs on rewarded and unrewarded trials, respectively. This is possible since rewarded trials will always elicit positive RPEs, and vice versa for unrewarded trials. The scaling of the firing rate as a function of, for example, positive prediction error is the regression weight used to *scale* the positive RPE in order to predict the firing rate, as in Dabney et al 2020. The size of the RPE is dependent on the cued probability at choice^8^. The RPE is defined as *r* − *V*, where *r* is the delivered reward and *V* is the cued probability value that denotes the expected value of the upcoming outcome. Rewarded and unrewarded trials yield a reward of 1 and 0, respectively. If, for example, the cued probability is high (0.9), this will elicit a smaller positive RPE on rewarded trials than a low cued probability (0.3), as reward was more expected (the RPEs in these cases would be: 1-0.9 = 0.1, and 1-0.3 = 0.7, respectively). In contrast, high cued probabilities will elicit larger negative RPEs on unrewarded trials, as reward was expected. We therefore estimate the scaling of positive and negative RPEs by regressing the chosen cue probability against the firing rate at feedback, separately for rewarded and unrewarded trials, resulting in the regression coefficients *β*^+^ and *β*^−^ for scalings of positive and negative RPEs, respectively. We use these scalings to compute the optimism of the scaling asymmetry as *β*^+^/(*β*^+^ + *β*^−^). These analyses are only carried out in RPE selective neurons, of which there were 41 in ACC (see below). The consistent diversity in reversal points observed at choice holds in this subset of neurons (Supplementary Figure 2). In order to confirm the revealed diversity is not simply due to noise, we perform the same partition-based consistency analysis as we did for optimism at choice.

This measure is analogous to the asymmetric scaling measure used in Dabney et al 2020^15^. The difference being that whereas Dabney et al 2020 measured asymmetric scaling from the cue presentation epoch, we estimate the scalings at a separate task epoch to cue presentation/choice; that is, at feedback time, when RPEs will be elicited following cued probabilistic reward delivery. Furthermore, we estimate positive and negative RPEs on rewarded and unrewarded trials, respectively.

To test for the relationship between optimism at choice and asymmetric scaling, we regressed choice optimism against asymmetric scaling across neurons. We perform this analysis for those neurons that encode RPEs. In other words, following Kennerley et al 2011^8^, those neurons that code the cued probability in choice epoch and feedback epoch with opposite signed relationships, and where both feedback epochs (rewarded and unrewarded) had the same sign. In brief, the logic is as follows: a RPE selective neuron that, for example, increases its firing rate as a function of chosen probability at choice, and therefore has a positive relationship between firing rate and RPE (elicited by the probability cue), should fire less strongly after reward following a high probability cue (since the RPE is smaller), and therefore has a negative relationship between firing rate between and probability at feedback. The same negative relationship between firing rate and probability at feedback applies on unrewarded trials, when a larger decrease in firing is elicited on high probability trials as a larger negative prediction error is elicited by lack of reward on high probability trials. Hence, the sign of the relationship between firing rate and probability cue is opposite at choice and feedback for RPE selective neurons as previous explained in detail^8^.

### Simultaneous diversity

Differences in value expectation may vary across sessions, due to, for example, motivation. Therefore, when pooling neurons across sessions for analysis, we might find diversity even from classic RL alone due to different expectations across sessions. To address this, we show that diversity exists within single sessions (Supplementary Figure 4). We further account for possible diversity across subjects in the asymmetry predicting reversal point correlation: we found that the relationship between choice reversal point and feedback asymmetric scaling held after including subject as a co-regressor (t(38) = 2.66; P=0.01, for the asymmetric scaling regressor predicting reversal point, in a GLM regressing out subject). This suggests differences in value expectations across subjects or sessions cannot explain the observed diversity.

### Task and neural recordings from Miranda et al (in prep): Two-step decision task

Results in Figure 3 are a re-analysis of the data presented in Miranda et al (in preparation), which are the neural recordings accompanying Miranda et al 2020^31^. Full task details can be found there, but we outline the key points relevant to the current study here.

This task was an adaptation of the classic two-step decision-making task^30^ to NHPs. The two-step nature of this task, along with the probabilistic transitions, is not relevant to the current study. This is because we focused analyses on the outcome time when learning of values of the stimuli ought to be occurring (see below). Nonetheless, we briefly describe the task here for completeness. Two decisions were made on each trial. At the first decision step, animals chose between two options (denoted by picture stimuli) that each result in probabilistic transitions to one of two second stage states. One transition was more likely (70% - a common transition) and the other less likely (30% - a rare transition). The common transition from each of the first stage options was to a different second stage option. In the each of the possible second stages, another two-option choice was required, and each of these 4 end-stage states had one of three different outcome levels (high, medium and low reinforcement levels) associated with it at a given time, which was delivered in the feedback stage. To induce learning, the outcome levels for the second stage options were dynamic: Reward associated with each second stage option remained the same for 5-9 trials, then changed randomly to any of the three possible outcome levels (including remaining the same). In order to make appropriate choices at both first and second stages of the task (which they did^31^), animals had to continually track and update the value of each end stage stimulus.

We focus exclusively on neural activity at the feedback stage when outcome was received. This is because: 1) we want to focus on the learning of the *dynamic* values of the second-stage options in order to test for asymmetric *learning*, 2) it is at this feedback period when RPEs ought to be elicited and error-driven learning of option values occurring, 3) this allows us to look at simple value learning without the added complication of the probabilistic transition structure of the task, which is not relevant for testing distributional RL. For the sake of our analyses, we can therefore think of this task as a simple reversal learning task in which four cues change their value every 5-9 trials. Amongst other brain regions, recordings were made in the ACC. We focus analyses on ACC since this is the brain region in common with Kennerley et al 2011 and where we have a strong hypothesis for the presence of distributional RL.

### Neurophysiological methods in the 2^nd^ (two-step) dataset

Two NHPs (Subjects “J” and “C”), different to those in Kennerley et al 2011, performed the task. Subjects were implanted with a titanium head positioner for restraint, then subsequently implanted with two recording chambers that were located on the basis of preoperative 3T MRI and stereotactic measurements. Postoperatively, we used gadolinium-attenuated MRI imaging and electrophysiological mapping of gyri and sulci to confirm chamber placement^18^. The chamber positioning along the anterior–posterior (AP), medial-lateral (ML) coordinate planes and their respective lateral tilt (LT) angle from vertical were as follows: one chamber over the left hemisphere at AP = 38(C)/37(J) mm, ML = 20.2(C)/18.1(J) mm and LT = 21◦(C)/26◦(J); and one over the right hemisphere at AP = 27(C)/27.5(J) mm, ML = 19.7(C)/17.9(J) mm and LT = 22.5◦(C)/28◦(J). Craniotomies were then performed inside each chamber to allow for neuronal recordings in different target regions.

For single-neuron recording we used epoxy-coated (FHC Instruments, Bowdoin, USA) or glass-coated (AlphaOmega Engineering, Nazareth, Israel) tungsten microelectrodes inserted through a stainless-steel guide tube mounted on a custom-designed plastic grid with 1 mm spacing between adjacent locations inside the recording chamber. Electrodes were acutely and slowly advanced through the intact dura at the beginning of every recording session using custom-built micro-drive assemblies manually controlled that lowered electrodes in pairs or triplets from a single screw; or motorised microdrives (Flex MT and EPS by Alpha Omega Engineering, Nazareth, Israel) with individual digital control of electrodes. During a typical recording session, 8–24 electrodes were lowered into multiple target regions until well-isolated neurons were found. Neuronal signals were acquired at 40 KHz, amplified, filtered and digitised (OmniPlex Neural Data Acquisition System by Plexon Instruments, Dallas, USA). Spike waveform sorting was performed off-line using principal component analysis-based method (Offline Sorter by Plexon Instruments, Dallas, USA). Channels were discarded if either neuronal waveforms could not be clearly separated, or if waveforms did not remain stable throughout the session.

We randomly sampled neurons; no attempt was made to select neurons on the basis of responsiveness or specific cortical layer. This procedure ensured an unbiased estimate of neuronal activity, thereby allowing a fair comparison of neuronal properties between the different brain regions.

We recorded neuronal data from the dorsal bank of anterior cingulate cortex (ACC). We used the gadolinium-enhanced MRI along with electrophysiological observations during the process of lowering each electrode to estimate the location of each recorded neuron. In ACC, the recordings were positioned between AP 30-37mm in Subject C, and AP 30-36mm in Subject J relative to the interaural line (AP=0mm).

### Neuron inclusion

Of the 240 neurons recorded in ACC, we tested for signatures of distributional RL (see below) in those that were sensitive to RPE (those neurons that had P<0.05 in linear regression between firing rate and RPE). The RPE regressors used to test for sensitivity are from Miranda et al 2020^31^. These are obtained using the best fitting parameters fit to behaviour, as described in Miranda et al 2020^31^. 111 neurons passed this criterion, and are the neurons analysed in Figure 3. Furthermore, we note that the results hold using a much more stringent definition of RPE from Bayer & Glimcher 2005^35^ (Supplementary Figure 5). That is, the firing rate at feedback on the current trial must be sensitive to the reward delivered on the current trial and on the previous trial, but with opposite signs, i.e. *FR* = *β*_0_ + *β*_1_*Rew*(*t*) + *β*_2_*Rew*(*t* − 1), where *β*_1_ and *β*_2_ are both significant at P<0.05 but with opposite signs. 39 neurons pass this criterion.

### Models and model fitting to test for asymmetric learning

#### Models

To test for asymmetric learning, we model neuron responses with classic and distributional RL models and test which is a better fit to the data. In all cases the model is used, for each neuron, to predict the firing rate on each trial (mean firing rate in a window of 200-600ms post feedback).

We adapt the one-step transition temporal difference learning model wherein estimates of cue values *V* are updated according to:

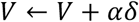

where *δ* is the reward prediction error, *δ* = *r* − *V*, where *r* is the reward delivered on the current trial and *V* the previous value estimate, and *α* is the learning rate by which *δ* is scaled to update values. This is the equation for classic RL and amounts to the Rescorla-Wagner model^10^.

The distributional RL version of this model is^15^:

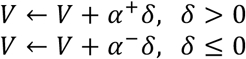

where *α*^+^ and *α*^−^ are separate learning rates for positive and negative RPEs/*δ*. In other words, the learning rate associated with a value update on a given trial will depend on whether the RPE was positive or negative. Different learning rates for positive and negative RPEs result in *asymmetries* in the rates at which neurons *learn* from better than expected and worse than expected feedback, i.e. *asymmetric learning*. This is unlike classic RL where learning is symmetric.

In order to fit the model to neural data, we predict the firing rate at feedback from the RPE. For the classic RL case this is as follows:

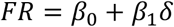

where *β*_0_ and *β*_1_ are regression coefficients. In the distributional RL case we have:

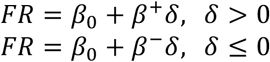

where *β*^+^ and *β*^−^ are different regression coefficients for positive and negative RPEs/*δ*, i.e. allows the FR to be a different *scaling* of the RPE for positive and negative RPEs. Critically, this *asymmetric scaling* is different to the above *asymmetric learning,* as it does not directly impact the update of the cue value *V*, and therefore subsequent computation of RPEs (*r* − *V*). It is therefore possible to have asymmetric scaling without asymmetric learning, and vice versa. Both asymmetric scaling and asymmetric learning are predictions of the distributional RL theory. Asymmetric scaling is what was tested for in Dabney et al 2020^15^ (it was not possible to test for learning in the task they analysed, nor the first dataset in this paper, due to the static nature of cue values).

We therefore have four possible models to test: symmetric scaling and symmetric learning (‘fully classic RL’), asymmetric scaling and symmetric learning, symmetric scaling and asymmetric learning, asymmetric scaling and asymmetric learning (‘fully distributional RL’).

#### Model fitting

We tested which of the above models were the best fit to the data. We did this by fitting the parameters in a subset of the data, and tested how well (measured using R-squared) a model using these fit parameters explained held out data in a 10-fold cross validation procedure. We then asked which model was the best fit to the data.

##### Fitting a simplified, single asymmetric scaling parameter

Since fitting all 4 parameters to the data was not possible due to computational demands, we adapt the asymmetric scaling equations such that asymmetric scaling can be accounted for with one, rather than two, parameters. We replace the asymmetric scaling equations with the following:

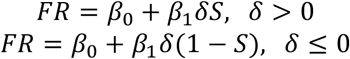

where *S* is bounded between 0 and 1 and acts as a single asymmetric scaling parameter (e.g. if *S* is near 1, positive RPE are scaled greatly relative to negative RPEs). Using *S* rather than fitting *β*^+^ and *β*^−^ therefore still achieves the important effect of accounting for asymmetries in the scaling of the FR by positive vs. negative RPEs. Note that the regression coefficients, *β*_0_ and *β*_1_, are the same in both equations, i.e. they were fit in the same regression model (positive and negative RPE trials were included in the same regression model), having scaled the RPEs by *S* or 1 − *S*. Note that it is the scaling parameter *S*, which captures the asymmetry, that is trained and tested in cross-validation, not the betas.

##### Estimating the parameters

We generate RPE regressors from each of the models and regress these against neural data. The regressors are generated by passing through the model the option chosen and reward observed on each trial of the training set. Values are updated and RPEs computed on each trial according to the above equations. We measure the model fit using the R-squared computed from the regression model. For each model (e.g. asymmetric learning with asymmetric scaling) we carry out the model fitting using a grid search over parameter space. Possible values for each parameter that is fit to the data – *α*^+^, *α*^−^ and *S* – lie between 0 and 1, and we perform the grid search with 0.025 size increments (this is an additional advantage to using *S* rather than *β*^+^ and *β*^−^, as the former but not the latter are bounded by 0 and 1, and can therefore be more easily fit with grid search). The combination of parameters with the highest R-squared is taken to be the best fit of parameters to the data.

##### Testing in held out data

For a given model, we take this combination of best-fitting parameters and use them to generate regressors in the held out data with the model equations above, again using the option chosen and reward delivered on each trial. We then assess their fit to the data by regressing the RPEs computed by the model in these held out trials against the firing rates on those trials. This results in R-squared values for the held out data, dependent on parameters fit to the training data, such that parameters capturing features of the data consistent across cross-validation folds will result in better fits in the held out data. We obtain 10 R-squared values for each model for each neuron, one for each cross-validation fold.

##### Testing for differences between model fits

We then compare, across the population of neurons, the different models’ fits to the neural data. We take the mean across the 10 cross-validated model fits in the test data for each model for each neuron, giving one number per model per neuron. We then carry out paired t-tests between the different models to determine the best fitting model. We found the asymmetric scaling with asymmetric learning model was better than all other models. This means the extra parameters improved the explanation of the neural data in the held out data (despite having to fit more parameters to the data), demonstrating that asymmetric learning is a better account of the data than symmetric learning.

## Acknowledgements

We would like to thank Philipp Schwartenbeck, Alon Baram, and Jacob Bakermans for very helpful discussions. S.V. was supported by the Leverhulme Doctoral Training Programme for the Ecological Study of the Brain. B.M. was supported by the Fundacão para a Ciência e Tecnologia (scholarship SFRH/BD/51711/2011) and the Prémio João Lobo Antunes 2017 - Santa Casa da Misericórdia de Lisboa. T.E.J.B. was supported by a Wellcome Trust Senior Research Fellowship (104765/Z/14/Z), Wellcome Trust Principle Research Fellowship (219525/Z/19/Z) together with funding from the James S McDonnell Foundation (JSMF220020372). S.W.K. was supported by NIMH (F32MH081521) and by Wellcome Trust Investigator Awards (096689/Z/11/Z and 220296/Z/20/Z).

## Supplementary Figures

**Supplemental Figure 1.**
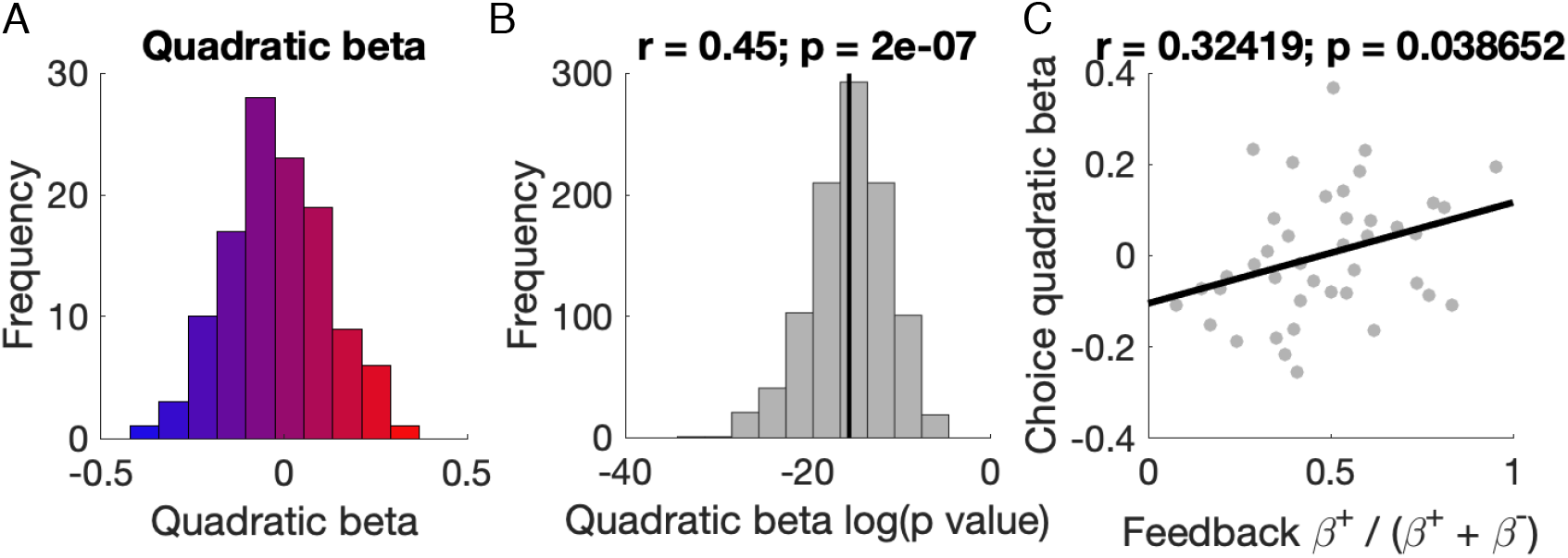
Results hold with a different measure of non-linearity at choice. A) Histogram showing diverse quadratic betas. B) Histogram showing the log p-values for consistency of these quadratic betas across partitions. C) Correlation between asymmetric scaling and quadratic betas.

**Supplemental Figure 2.**
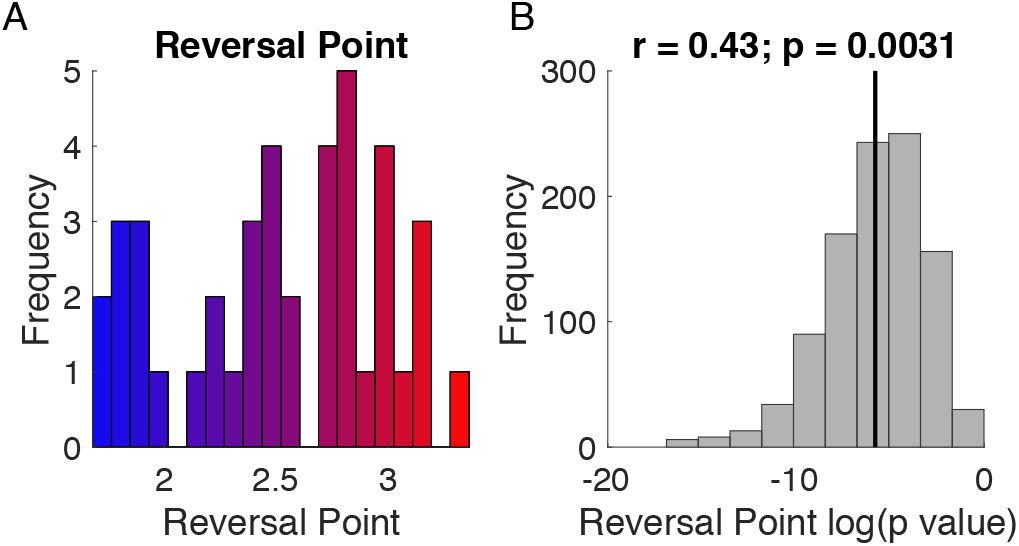
Consistent and diverse reversal points at choice in only those neurons defined as RPE selective. Same analyses as in Figure 1 testing for consistent and diverse reversal points, but in only those neurons that are RPE selective, i.e. those neurons in Figure 2 (as opposed to all reward-sensitive neurons, as in Figure 1). A) Diversity in reversal points. B) This diversity is consistent.

**Supplemental Figure 3.**
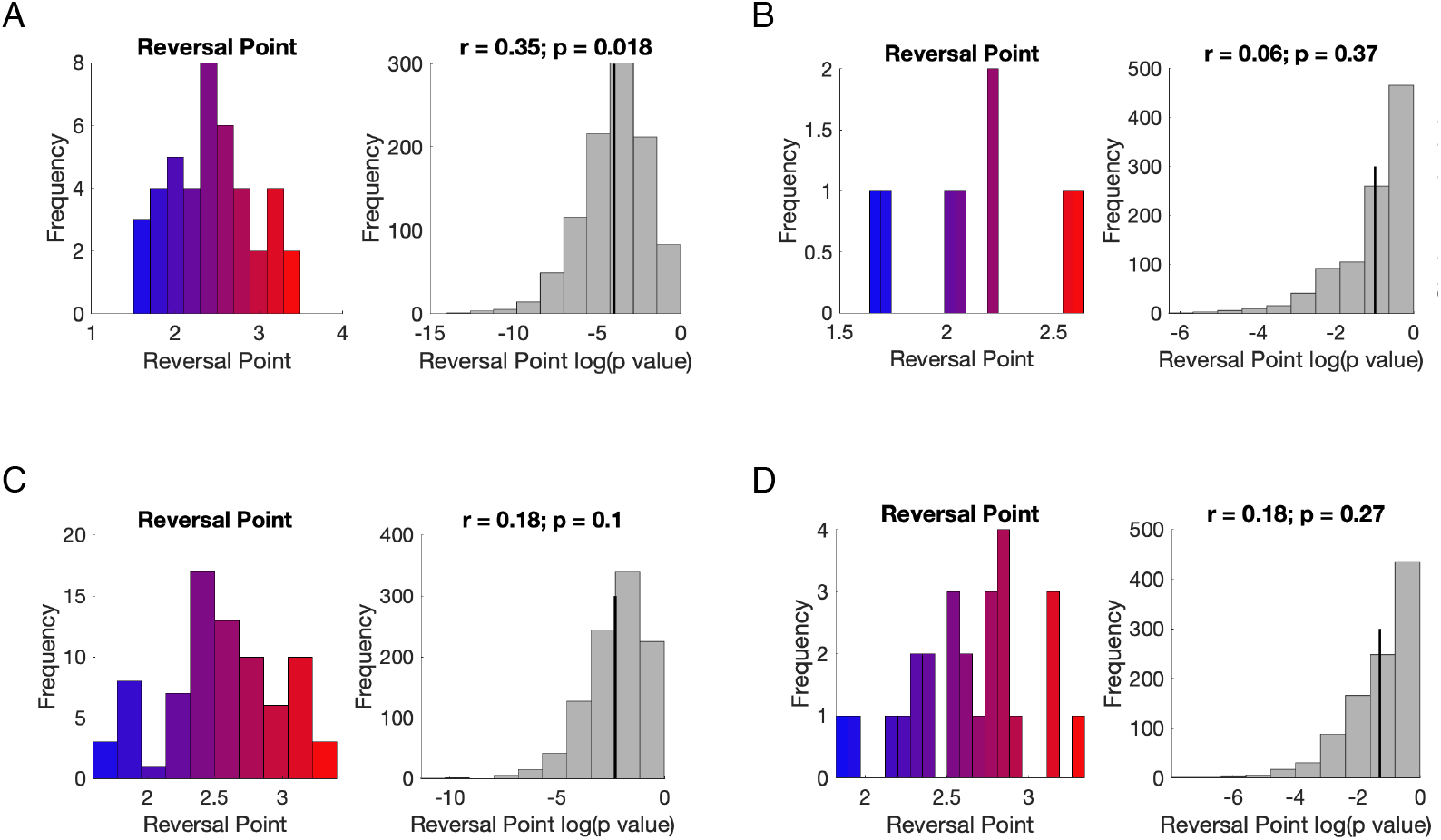
Lack of diversity in OFC and LPFC. Same analysis as in Figure 1 and Supplemental Figure 2, but for OFC and LPFC on all reward-selective neurons or RPE selective neurons. A) OFC reward-selective neurons. B) OFC RPE-selective neurons. C) LPFC reward-selective neurons. D) LPFC RPE-selective neurons. With the exception of the reward-selective neurons in OFC (A), none of these analyses were significant. The RPE-selective neurons had no consistent diversity, so we did not look for further distributional RL signatures in these brain regions.

**Supplemental Figure 4.**
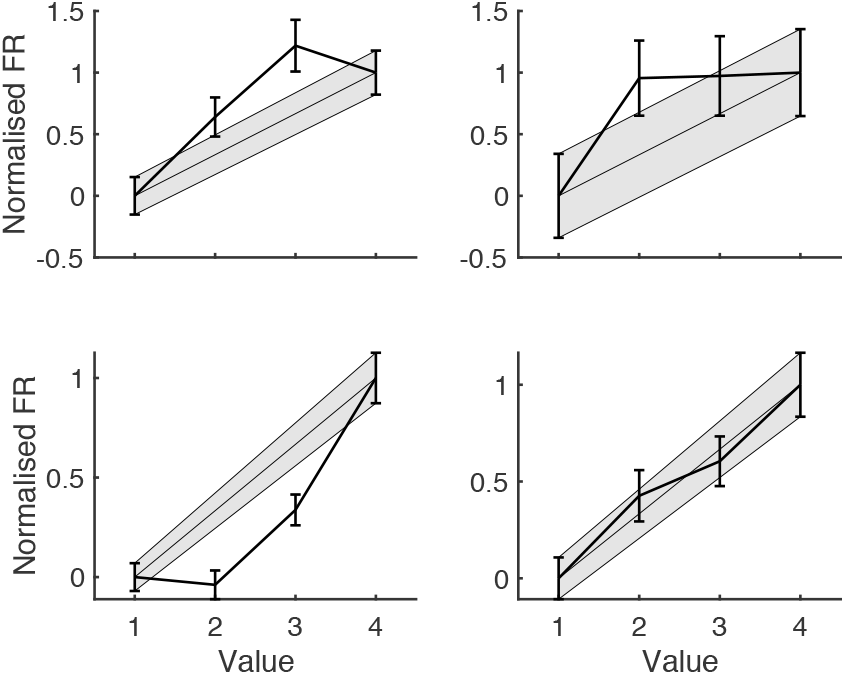
Simultaneous diversity within session. Four simultaneously recorded cells from the session with most reward-sensitive cells (9 in total), demonstrates there is diversity in optimism even within a session. Across cells, responses to middle value levels are both above and below the linear interpolation between lowest and highest values’ responses. Mean normalised firing is plotted for each of the 4 value levels. Firing rates are normalised such that responses to value 1 and 4 have mean firing rate 0 and 1, respectively. Normalisation allows comparison across cells of responses to middle value levels. Responses to value 2 across the 9 simultaneously recorded cells were significantly diverse; ANOVA rejected the null hypothesis that across cells the value 2 responses were drawn from the same mean (*F*(8,405) = 3.56, *P* = 0.0005). The same was true for responses to value 3 (*F*(8,441) = 2.16, *P* = 0.0291).

**Supplementary Figure 5.**
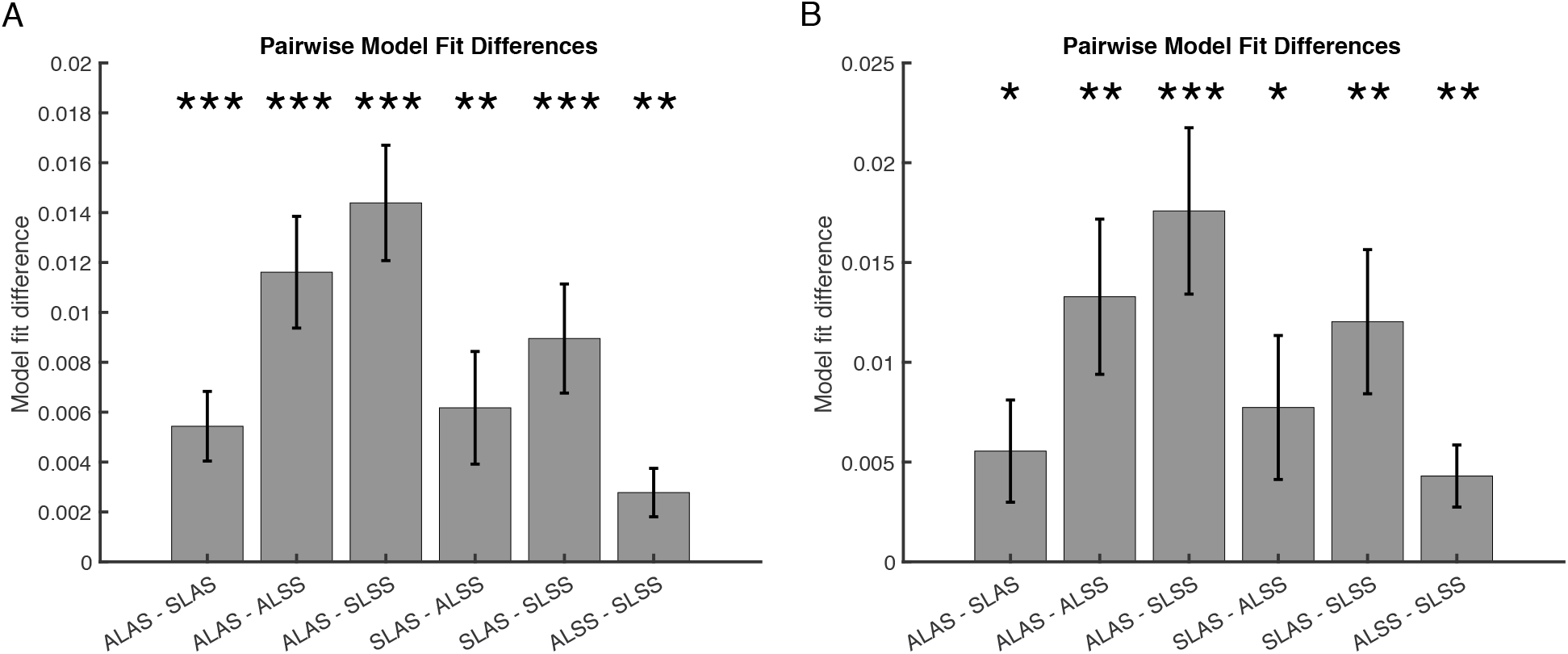
All pairwise model comparisons for asymmetric learning and scaling analyses. Same format as Figure 3 in the main text. A) Bar graphs with all 6 pairwise model comparisons. ALAS – SLAS, SLAS – ALSS, and ALSS – SLSS are the same as in the main text. B) Same as A but for only those neurons (n=39) that meet a strict definition for being RPE selective. That is, as defined in Bayer & Glimcher 2005, those neurons that encode reward on the current trial and previous trial but with opposite signs.

